## Supplementary Information for "DeepRTAlign: toward accurate retention time alignment for large cohort mass spectrometry data analysis"

### Supplementary Methods

#### Proteomics sample preparation.

All the HCC tissues in cohort HCC-R2 were obtained from the Eastern Hepatobiliary Surgery Hospital, Naval Medical University, Shanghai, China. All experiments were approved by the Research Ethical Committee of Eastern Hepatobiliary Surgery Hospital. Filter-aided sample preparation (FASP) was employed for tissue sample pretreatment. A 500 μg aliquot of liver proteins was diluted to 500 μL with a UA solution containing 8 M urea in 100 mM Tris-HCl (pH 8.5). Following centrifugation on a 30-kDa filter for 20 min, 200 μL of UA solution containing 10 mM DTT was added and the reaction was allowed to proceed at 37 °C for 4 h. After removal of the solution, a UA solution containing 50 mM iodoacetamide (IAA) was introduced and let react in the dark at room temperature (RT) for 30 min. The ultra-fraction tubes were then washed thoroughly with 200 μL of UA three times and 200 μL of 50 mM ammonium bicarbonate (ABC) three times. Next, 100 μL of ABC containing 0.1 μg/μL trypsin was added and let incubate at 37 °C for 12 h. The filter tubes were then washed twice with 100 μL of ABC via centrifugation, and flow-through fractions were pooled. Peptide concentration was determined using a NanoDrop One at 280 nm. Peptide mixtures were dried in a SpeedVac and stored at -80 °C before use.

Human embryonic kidney (HEK) 293T cells were cultured and harvested as previously reported[1]. *E. coli* (DH5a) cell lysate was purchased from MCLAB (ECCL-100). Both 293T cells and *E. coli* cell lysate were then treated by the same in-solution digestion method to obtain peptides. First, lysis buffer containing 10% Sodium deoxycholate (DOC), 1 M Tris HCl (pH=8.8), 100 mM Tris (2-carboxyethyl) phosphine (TCEP) and 400 mM 2-Chloroacetamide (CAA) was added to the solutions. Solutions were heated at 95°C for 5 min on a ThermoMixer (Eppendorf) and cooled down on ice for 5 min. Next, solutions were sonicated (Ningbo scientz, 25% energy, 30 s) on ice-water. Subsequently, proteins were digested overnight with 1:25 (w/w) MS-grade trypsin (Promega, V5280) at 37 °C. Finally, digested peptides were desalted using C18 solid-phase extraction (SPE) column (Waters), dried in SpeedVac, and stored at -20 °C before nanoLC-MS analysis. The concentration of the peptides was determined using a NanoDrop One at a wavelength of 280 nm.

#### Targeted LC-MS/MS analysis (dataset HCC-R2)

NanoLC-MS experiments were performed using Vanquish Neo system interfaced with an Orbitrap Exploris 480 mass spectrometer (Thermo Fisher Scientific, USA) operated in scheduled parallel-reaction-monitoring (sPRM). Peptide samples were separated on homemade capillary columns (100 μm i.d. x 30 cm) with integrated spray tips. The columns were packed with 1.9 μm/120 Å ReproSil-Pur C18 resins (Dr. Maisch GmbH, Germany), and all columns were heated at 55 °C. In sPRM mode, 500 ng of HCC peptide sample was injected each time, and a bit 1x iRT kit (Biognosys) was spiked in each sample for retention time calibration in Skyline analysis.

Mobile phases A and B were water and 80%/20% acetonitrile (ACN)/water (v/v) with 0.1% FA (v/v), respectively. The flow rate was 300 nL/min across the total gradients. The segmented 190-min gradient used in PRM mode is the following: 3%-8% (v/v) buffer B in 4.5 min, 8%-30% (v/v) buffer B for 168 min, 30%-40% (v/v) buffer B for 7.5 min, followed by a 2 min wash from 40% to

95% (v/v) buffer B. In the end, 95% (v/v) buffer B was kept for 8 min. After the gradient, a 20 min wash from 3% to 95% (v/v) buffer B was performed to minimize potential carryover. Scheduled PRM parameters were as follows: Full scan (MS1) from m/z 335 to 1 120 was acquired at the resolution of 120 000. The automatic gain control (AGC) target was 1E6, and the maximum injection time (maxIT) was set to 200 ms. A subset of the top 200 MS features, the 15 features and other 34 features (a total of 49 features), were selected as targeted precursors in the mass list table (**Supplementary Table 15**) for further MS/MS scans. Targeted precursors were isolated through a window of 1.2 Th. The MS2 scans were acquired at a resolution of 120 000 with an AGC setting of 4E5, a maxIT of 200 ms, and a normalized collision energy (NCE) of 28%.

**Nano-flow LC-MS/MS analysis of HEK 293T and *E. coli* mixtures (dataset Benchmark-FC)**

The Vanquish Neo system, Orbitrap Exploris 480 mass spectrometer, mobile phases, and capillary columns were the same as described above for targeted LC-MS/MS analysis. The flow rate was 300 nL/min, except that 500 nL/min was applied in the first 1.5 min of the total gradient. The gradient used in nano-flow LC-MS/MS analysis of 293T and *E.coli* mixtures is: 3%-8% (v/v) buffer B in 1.5 min, 8%-30% (v/v) buffer B for 55 min, 30%-40% (v/v) buffer B for 3.5 min, followed by a 1.5 min wash from 40% to 95% (v/v) buffer B, and 95% (v/v) buffer B was kept for 18.5 min at the end. DDA parameters were as follows: Full MS scans over the m/z range of 350-1 500 were performed at a resolution of 60 000. The AGC target was set to 3E6, and the maximum injection time was 45 ms. MS/MS acquisition was performed in top speed mode with a 1.5-s cycle time. The resolved fragments were scanned at a mass resolution of 15 000 and an AGC target value of 3E5. The threshold to trigger MS2 scans was 5E3, and the maxIT was 22 ms. Ions with charge states of 2-6 were sequentially fragmented by higher-energy collision dissociation (HCD) with an NCE of 28%. The isolation window was 1.6 Th, and the dynamic exclusion time was set to 40 s. 200 ng of 293T peptides spiked with 10, 15, 20, or 25 ng of *E. coli* peptides was loaded in each run, and 0.5 µL of 2.5x iRT kit was spiked in each sample. Each group contains three technical replicates.

**Micro-flow LC-MS/MS analysis of pure HEK 293T or *E. coli* (datasets Benchmark-QC-H and Benchmark-QC-E)**

A Dionex UltiMate 3000 Rapid Separation LC (RSLC) system was online coupled to an Orbitrap Exploris 480 mass spectrometer (Thermo Fisher Scientific) with an OptaMax NG Atmospheric Pressure Ionization Source (H-ESI mode). Commercially available Thermo Fisher Scientific Acclaim PepMap 100 C18 LC columns (1 mm ID × 150 mm, 2 µm particle size, catalog number 164711) were used for peptide separations. The column temperature was maintained at 55 °C using the integrated column oven in the LC system. Flow rate of 50 µL/min was used to deliver the segmented gradient, and mobile phases A and B were water and ACN with 0.1% FA (v/v), respectively. The segmented gradient is: 0.5%-5% (v/v) buffer B in 0.5 min, 5%-32% (v/v) buffer B for 60 min, 32%-95% (v/v) buffer B for 0.2 min. In the end, 95% (v/v) buffer B was kept for 2.5 min. After the gradient, columns were equilibrated 0.5% B for 1.8 min before the next injection. DDA data were acquired using the following parameters: Full scans were acquired in Orbitrap at a resolution of 60 000 (m/z 200) and AGC value of 3E6. The isolation window was 1.3 Th and m/z range was 350-1 550 in full scans. The top 30 precursors found in full scans were selected for fragmentation. MS/MS scans were acquired with 28% normalized collision energy in HCD mode at a resolution of 15 000 (m/z 200), charge states of 2-6 and minimum intensity of 1 000. AGC target value for fragment spectra was set to 1.5E5. For MS2 spectra, the maxIT was set to 30 ms, and dynamic exclusion time was set to 30 s. 10 µg of 293T or *E. coli* tryptic peptides were loaded each time and the DDA data were collected from three Orbitrap Exploris 480 mass spectrometers with the same settings.

**Nano-flow LC-MS/MS analysis of 293T digest (dataset Benchmark-RT)**

The Dionex UltiMate 3000 system, Orbitrap Exploris 480 mass spectrometer and mobile phases, were the same as described above for micro-flow LC-MS/MS analysis but operated in nano-flow mode. DDA parameters were the same as described in Nano-flow LC-MS/MS analysis of 293T and *E. coli* mixtures except that NCE was set to 30% instead of 28%. Homemade capillary columns (100 μm i.d. x 20 cm) with integrated spray tips were applied for LC separation. Two different gradients (60 and 120 min active gradients) were used with flow rate of 250 nL/min. Samples were loaded into capillary columns in 15 min at 700 nL/min, followed by flow rate decreased to 250 nL/min in 4.5min. The active 60 min gradient is: 0.5%-6.4% (v/v) buffer B in 2 min, 6.4%-24% (v/v) buffer B for 55 min, 24%-32% (v/v) buffer B for 3.5 min. In the end, 32% (v/v) buffer B was increased to 95% in 1.5min and kept for 5 min. The active 120 min gradient is the same as the active 60 min gradient, except that 6.4% buffer B increased to 24% (v/v) in 115 min. 1 μg of 293T digest was injected in each run.

**PRM data analysis**

PRM raw data were firstly searched against the human UniProt FASTA database (downloaded on September 27, 2018) using Proteome Discoverer (PD) 2.5 software with the Sequest HT search engine to identify peptide sequences. Trypsin was selected as the proteolytic enzyme, and missed cleavage sites were allowed up to two. Cysteine carbamidomethylation was set as the static modification. The oxidation of M and acetylation of the protein N-terminal were set as the dynamic modifications. The precursor mass tolerance was set to 10 ppm, and the fragment mass tolerance was 0.02 Da. The false discovery rates of the peptide-spectrum matches (PSMs) and proteins were set to less than 1%.

PRM data files were further analyzed using Skyline 22.2 (MacCoss Lab)[3]. A merged spectral library was generated in Skyline from MSF files of PD results and used as the reference library. An iRT database was generated along with the merged library for RT prediction, and peptide peaks were filtered to within 5 min of the predicted RT. The digestion enzyme was set to trypsin. Precursor charges 2 to 7 and ion charges 1 to 3 were allowed. Ion match tolerance was set to 0.02 Th for the selection of fragment ions from the spectral library. Raw data files were imported into Skyline for automated peak detection using "targeted" MS/MS filtering mode with the mass analyzer set to Orbitrap. For those identified 52 precursors from 48 MS features, auto-picked peaks in Skyline were further filtered manually to fulfill the following criteria: dotp values >0.7, mass error within 5 ppm, detection of at least 5 fragments, and consistent RT across all 23 HCC samples. Peptides in a few samples that did not meet these criteria were considered undetected, and their peak areas in those samples were assigned a value of 0. Peptide abundance values of the 15 features were exported from Skyline. Peptide abundance was obtained by summing the peak area of both precursor ions and selected fragment ions.

**DDA data analysis**

MSFragger v3.7 (in FragPipe v19.1), Mascot v2.8.1, MaxQuant v1.6, and PD 2.5 were used for DDA data analysis. HCC-T, HCC-N and HCC-R datasets were searched against the human UniProt FASTA database (downloaded on September 27, 2018). UPS2-M dataset was searched against the mouse UniProt FASTA database (downloaded on November 4, 2015). UPS2-Y dataset was searched against the yeast UniProt FASTA database (downloaded on June 15, 2021). EC-H dataset was searched against the FASTA database provided in PRIDE (https://www.ebi.ac.uk/pride/archive/projects/PXD003881). AT dataset was searched against the mouse-ear cress UniProt FASTA database (downloaded on August 2, 2021).

In MSFragger, the digestion enzyme was set to trypsin. Precursor mass tolerance and fragment mass tolerance were 20 ppm, peptide length was limited to 7-50. In MaxQuant, precursor mass tolerance and fragment mass tolerance are 20 ppm, min peptide length is 7. In Mascot, precursor mass tolerance is 20 ppm, and fragment mass tolerance is 0.05 Da. For database searching of 293T DDA data collected in two gradients, PD 2.5 settings are the same as used for sPRM data. For all the tools, fixed and variable modifications, digestion enzyme, and FDR rate were the same as described above for PD 2.5 settings of sPRM data. The other parameters were set to default in these tools.

**DIA data analysis**

DIA-NN v1.8 was used for DIA data analysis. Dataset SC were searched against the FASTA database provided in PRIDE (https://www.ebi.ac.uk/pride/archive/projects/PXD025634). The digestion enzyme was set to trypsin, missed cleavage was 1, peptide length range was 7-30, and precursor charge range was 1-4. Cysteine carbamidomethylation was set as the fixed modification, while the protein N-terminal were set as the variable modification. The other parameters were the default settings in DIA-NN.

### Supplementary Tables

**Supplementary Table 1.** The datasets used for training and testing the deep learning model of DeepRTAlign in this study. The sample numbers in this table were the number of samples used in this work. HCC-T and HCC-N indicated the data from tumor and non-tumor samples of an HCC cohort (N=101). HCC-R and HCC-R2 were data from two HCC cohorts. UPS2-M and UPS2-Y were two benchmark datasets from mouse cells and yeast cells with UPS2 proteins spiked in. EC-H was a dataset from the mixture of human cells and *E. coli* cells. AT was a dataset based on the *Arabidopsis thaliana* seeds. SC was a single-cell proteomic dataset. MI was based on mouse intestinal samples. CD was obtained from the gut microbiota of patients with Crohn's disease. NCC19, SM1100, MM, SO and GUS were public metabolomic datasets. Benchmark-QC-H and Benchmark-QC-E were two benchmark datasets based on HEK 293T and *E. coli* samples, respectively. Benchmark-FC was a benchmark dataset with known fold changes. Benchmark-RT contained two HEK 293T samples with different RT gradients (60 min and 120 min).

| Dataset name | Sample numbers | Dataset ID | RT range (min) | Type |
|---|---|---|---|---|
| HCC-T | 101 | PXD006512 | 80 | Training set |
| HCC-N | 101 | PXD006512 | 80 | Proteomic test set |
| HCC-R | 11 | PXD022881 | 60 | Proteomic test set |
| UPS2-M | 12 | PXD008428 | 100 | Proteomic test set |
| UPS2-Y | 12 | PXD008428 | 100 | Proteomic test set |
| EC-H | 20 | PXD003881 | 170 | Proteomic test set |
| AT | 18 | PXD027546 | 130 | Proteomic test set |
| SC | 18 | PXD025634 | 90 | Proteomic test set |
| MI | 1 | PXD002838 | 180 | Proteomic test set |
| CD | 1 | PXD002882 | 120 | Proteomic test set |
| NCC19 | 1 | MTBLS1866 | 30 | Metabolomic test set |
| SM1100 | 10 | MTBLS733 | 50 | Metabolomic test set |
| MM | 1 | MTBLS5430 | 40 | Metabolomic test set |
| SO | 1 | MTBLS492 | 45 | Metabolomic test set |
| GUS | 1 | MTBLS650 | 40 | Metabolomic test set |
| HCC-R2 | 23 | IPX0006622000 | 180 | PRM validation |
| Benchmark-FC | 12 | IPX0006638000 | 60 | Benchmark (known fold changes) |
| Benchmark-QC-H | 3 | IPX0006819000 | 60 | Benchmark for QC |
| Benchmark-QC-E | 3 | IPX0006819000 | 60 | Benchmark for QC |
| Benchmark-RT | 2 | IPX0006820000 | 60 and 120 | Alignment for different gradients |

**Supplementary Table 2.** Parameters optimization for the DNN model in DeepRTAlign based on the 10-fold cross validation results of the training set HCC-T.

(a) Optimization for hidden layer number in the DNN model. In this test, each layer has 5000 neurons.

| Hidden layer number | 1 | 2 | 3 | 4 | 5 |
|---|---|---|---|---|---|
| AUC | 0.988±0.003 | 0.990±0.002 | **0.993±0.002** | 0.992±0.003 | 0.993±0.002 |

(b) Optimization for neuron number in the DNN model. All the models have 3 hidden layers.

| Neuron number | 50 | 500 | 5000 | 50000 | 500000 |
|---|---|---|---|---|---|
| AUC | 0.887±0.012 | 0.969±0.011 | **0.993±0.002** | 0.993±0.001 | 0.992±0.001 |

**Supplementary Table 3.** The AUCs on different test sets. All the results are based on the model trained on the HCC-T dataset. In each test set, we randomly selected 10,000 positive and 10,000 negative feature pairs to perform this evaluation.

| Dataset | DNN | RF | KNN | SVM | LR |
|---|---|---|---|---|---|
| HCC-N | **0.925** | 0.916 | 0.656 | 0.865 | 0.894 |
| HCC-R | **0.933** | 0.905 | 0.668 | 0.901 | 0.899 |
| UPS2-M | **0.979** | 0.919 | 0.683 | 0.896 | 0.905 |
| UPS2-Y | **0.971** | 0.920 | 0.702 | 0.900 | 0.897 |
| EC-H | **0.972** | 0.938 | 0.733 | 0.912 | 0.944 |
| AT | **0.975** | 0.943 | 0.785 | 0.932 | 0.945 |
| SC | **0.917** | 0.901 | 0.752 | 0.842 | 0.898 |

**Supplementary Table 4.** Parameters optimization for the RF model based on the 10-fold cross validation results of the training set HCC-T.

(a) Optimization for n_estimators in the RF model. All the other parameters were kept default in scikit-learn v0.21.3.

| n_estimators | 10 | 50 | 100 | 150 |
|---|---|---|---|---|
| AUC | 0.964±0.030 | **0.974±0.025** | 0.973±0.026 | 0.974±0.025 |

(b) Optimization for max_depth in the RF model. n_estimators was set to 50. All the other parameters were kept default in scikit-learn v0.21.3.

| max_depth | 10 | 50 | 100 | None |
|---|---|---|---|---|
| AUC | 0.966±0.025 | **0.974±0.026** | 0.973±0.027 | 0.973±0.024 |

**Supplementary Table 5.** Parameters optimization for K in the KNN model based on the 10-fold cross validation results of the training set HCC-T. All the other parameters were kept default in scikit-learn v0.21.3.

| K | 1 | 2 | 3 | 4 | 5 | 6 |
|---|---|---|---|---|---|---|

| | | | | | | |
|---|---|---|---|---|---|---|
| AUC | 0.807±0.080 | 0.836±0.085 | 0.850±0.083 | 0.853±0.083 | **0.853±0.081** | 0.852±0.077 |

**Supplementary Table 6.** Parameters optimization in the SVM model based on the 10-fold cross validation results of the training set HCC-T. All the other parameters were kept default in scikit-learn v0.21.3.

(a) Optimization for kernel in the SVM model. All the other parameters were kept default in scikit-learn v0.21.3.

| kernel | "linear" | "poly" | "rbf" | "sigmoid" |
|---|---|---|---|---|
| AUC | 0.908±0.018 | **0.910±0.034** | 0.890±0.054 | 0.637±0.090 |

(b) Optimization for gamma in the SVM model. The kernel was set to "poly". All the other parameters were kept default in scikit-learn v0.21.3.

| gamma | "scale" | "auto" | 0.01 | 0.1 | 1 |
|---|---|---|---|---|---|
| AUC | 0.860±0.022 | **0.911±0.034** | 0.812±0.013 | 0.902±0.032 | 0.907±0.031 |

(c) Optimization for C in the SVM model. The kernel was set to "poly". Gamma was set to "auto". All the other parameters were kept default in scikit-learn v0.21.3.

| C | 0.1 | 1 | 5 |
|---|---|---|---|
| AUC | 0.903±0.028 | **0.910±0.034** | 0.901±0.030 |

(d) Optimization for degree in the SVM model. The kernel was set to "poly". Gamma was set to "auto" and C was set to 1. All the other parameters were kept default in scikit-learn v0.21.3.

| degree | 1 | 3 | 5 |
|---|---|---|---|
| AUC | 0.901±0.018 | **0.910±0.033** | 0.904±0.025 |

**Supplementary Table 7.** Parameters optimization in the LR model based on the 10-fold cross validation results of the training set HCC-T. All the other parameters were kept default in scikit-learn v0.21.3.

(a) Optimization for solver in the LR model. All the other parameters were kept default in scikit-learn v0.21.3.

| solver | lbfgs | liblinear | newton-cg | sag | saga |
|---|---|---|---|---|---|
| AUC | **0.912±0.018** | 0.911±0.017 | 0.911±0.017 | 0.911±0.017 | 0.911±0.017 |

(b) Optimization for penalty in the LR model. The solver was set to "lbfgs". All the other parameters were kept default in scikit-learn v0.21.3.

| penalty | L2 | None |
|---|---|---|
| AUC | **0.911±0.017** | 0.911±0.017 |

**Supplementary Table 8.** The minimum information required for alignment in each tool. Symbol "√" represents for required and "-" represents for "not required".

| Tools | MS | MS/MS | Identification results |
|---|---|---|---|
| DeepRTAlign | √ | - | - |
| MZmine 2 | √ | - | - |
| OpenMS | √ | - | - |
| Quandenser | √ | √ | - |
| MaxQuant | √ | √ | √ |
| MSFragger | √ | √ | √ |
| DIA-NN | √ | √ | √ |

**Supplementary Table 9.** The AUCs of DeepRTAlign when using different samples in the test sets as the anchor sample. All the results are based on the model trained on the HCC-T dataset. In each test set, five samples are randomly selected.

| Dataset | Sample 1 | Sample 2 | Sample 3 | Sample 4 | Sample 5 |
|---|---|---|---|---|---|
| HCC-N | 0.925 | 0.926 | 0.925 | 0.926 | 0.924 |
| HCC-R | 0.933 | 0.930 | 0.930 | 0.933 | 0.934 |
| UPS2-M | 0.979 | 0.977 | 0.976 | 0.976 | 0.981 |
| UPS2-Y | 0.971 | 0.972 | 0.973 | 0.971 | 0.971 |
| EC-H | 0.972 | 0.971 | 0.972 | 0.972 | 0.972 |
| AT | 0.975 | 0.975 | 0.974 | 0.976 | 0.975 |
| SC | 0.917 | 0.909 | 0.915 | 0.919 | 0.918 |

**Supplementary Table 10.** The AUCs of DeepRTAlign with or without coarse alignment step in different test sets. All the results are based on the model trained on HCC-T dataset. And in this table, all the models have 3 hidden layers, and each layer has 5000 neurons.

| Dataset | With coarse alignment | Without coarse alignment |
|---|---|---|
| HCC-N | **0.925** | 0.899 |
| HCC-R | **0.933** | 0.875 |
| UPS2-M | **0.979** | 0.909 |
| UPS2-Y | **0.971** | 0.898 |
| EC-H | **0.972** | 0.905 |
| AT | **0.975** | 0.917 |
| SC | **0.917** | 0.821 |

**Supplementary Table 11.** The AUCs of DeepRTAlign when using random features to replace the original features in the test sets. All the results are based on the model trained on the HCC-T dataset. “Difference” means the differences of RT and m/z in the samples to be aligned (i.e., the part 2 and part 3 in **Figure 1a**). “Original” means the original RT and m/z values in the samples to be aligned (i.e., the part 1 and part 4 in **Figure 1a**). “All” means all the features used in DeepRTAlign shown in **Figure 1a**.

| Dataset | m/z | RT | Difference | Original | All |
|---|---|---|---|---|---|
| HCC-N | 0.899 | 0.841 | 0.912 | 0.924 | 0.499 |
| HCC-R | 0.895 | 0.855 | 0.905 | 0.930 | 0.502 |
| UPS2-M | 0.952 | 0.932 | 0.923 | 0.979 | 0.499 |
| UPS2-Y | 0.958 | 0.941 | 0.946 | 0.970 | 0.499 |
| EC-H | 0.959 | 0.942 | 0.938 | 0.970 | 0.512 |
| AT | 0.964 | 0.950 | 0.945 | 0.972 | 0.498 |
| SC | 0.901 | 0.831 | 0.844 | 0.917 | 0.503 |

**Supplementary Table 12**. The different algorithm combinations for benchmarking DeepRTAlign against MZmine 2 and OpenMS on a public metabolomic test set SM1100.

| Abbreviations | Feature extraction | Feature alignment | Precision | Recall |
|---|---|---|---|---|
| MM | MZmine 2 | MZmine 2 | 1.000 | 1.000 |
| MD | MZmine 2 | DeepRTAlign | 1.000 | 1.000 |
| OO | OpenMS | OpenMS | 1.000 | 0.980 |
| OD | OpenMS | DeepRTAlign | 0.997 | 0.985 |
| DD | Dinosaur | DeepRTAlign | 0.971 | 0.965 |

**Supplementary Table 13.** The 15 aligned MS1 features in the feature-based early-recurrence classifier for HCC early recurrence prediction. Their identification results from both DDA data and sPRM data were provided. This table is presented as an Excel file.

**Supplementary Table 14.** Clinical information of the HCC-R2 cohort (N=23). This table is presented as a csv file.

**Supplementary Table 15. The mass list of target precursors for PRM experiment.** A total of 49 features including the 15 features and other 34 features, were selected from the top 200 MS features in dataset HCC-T, as the targeted precursors in the mass list table in the PRM experiment. 11 iRT standards (iRT kit, Biognosys) was included in the mass list for retention time calibration in Skyline analysis. For four features, precursors with two different charges were included in the mass list. Therefore, a total of 64 targeted precursors were set in the mass list. Targeted precursors were isolated through a window of 1.2 Th. Normalized collision energy was set to 28%. This table is presented as a csv file.

**Supplementary Table 16.** The peptide identification results of PD 2.5 for the PRM data of HCC-R2 cohort (N=23). Trypsin was selected as the proteolytic enzyme, and missed cleavage sites were allowed up to two. Cysteine carbamidomethylation was set as the static modification. The oxidation of methionine and the acetylation of the protein N-terminal were set as the dynamic modifications. The precursor mass tolerance was 10 ppm, and the fragment mass tolerance was 0.02 Da. The false discovery rates of the peptide-spectrum matches (PSMs) and proteins were set to less than 1%. This table is presented as an Excel file.

**Supplementary Table 17.** Quantification results of the verified 48 features exported from Skyline, including precursors and transitions. Peak areas of precursors and transitions were summed as peptide abundance of the 15 features and used for the prediction of early recurrence. This table is presented as a csv file.

**Supplementary Table 18.** The AUCs of different DNN hidden layer numbers of Siamese network on different test sets. All the results are based on the model trained on HCC-T dataset. And in this test, each layer has 5000 neurons.

| Dataset | 1 hidden layer | 2 hidden layers | 3 hidden layers | 4 hidden layers | 5 hidden layers |
|---|---|---|---|---|---|
| HCC-N | 0.725 | 0.719 | 0.799 | **0.802** | 0.743 |
| HCC-R | 0.759 | 0.771 | **0.789** | 0.774 | 0.781 |
| UPS2-M | 0.713 | 0.702 | 0.754 | 0.761 | **0.762** |
| UPS2-Y | 0.771 | **0.792** | 0.705 | 0.734 | 0.755 |
| EC-H | 0.723 | 0.773 | **0.815** | 0.797 | 0.798 |
| AT | 0.748 | 0.744 | 0.735 | **0.781** | 0.712 |
| SC | **0.733** | 0.713 | 0.713 | 0.720 | 0.716 |

**Supplementary File 1.** Skyline annotated spectra for the verified 48 features using PRM.

### Supplementary Figures

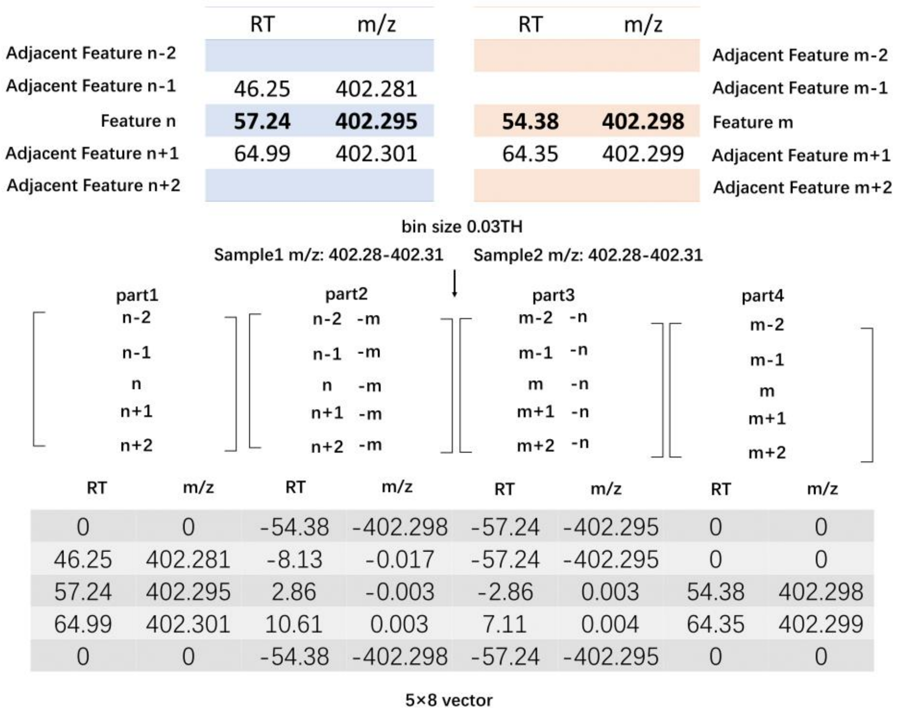


**Supplementary Fig. 1.** An input example for DeepRTAlign. After min-max normalization on each column, this 5×8 vector is used as the input to the neural network. If feature n and feature m are the same peptide, this vector will be labeled as "aligned" (should be aligned), otherwise it will be labeled as "non-aligned" (should not be aligned).

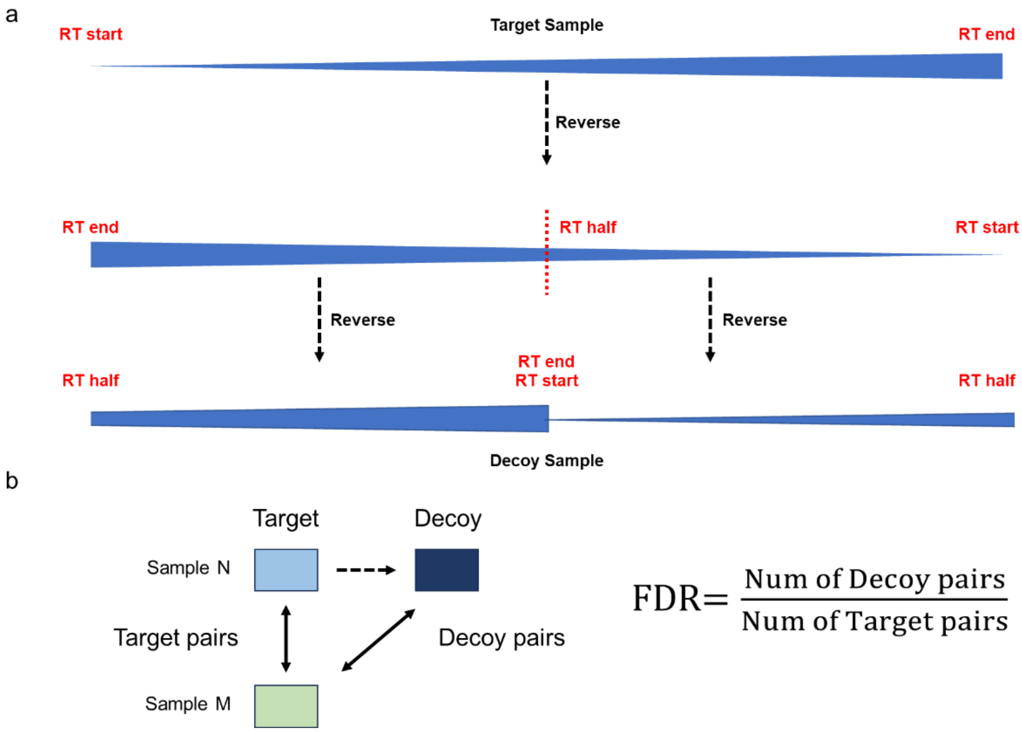


**Supplementary Fig. 2.** Illustration of the QC module in DeepRTAlign. (a) The decoy design workflow. (b) The FDR calculation workflow.

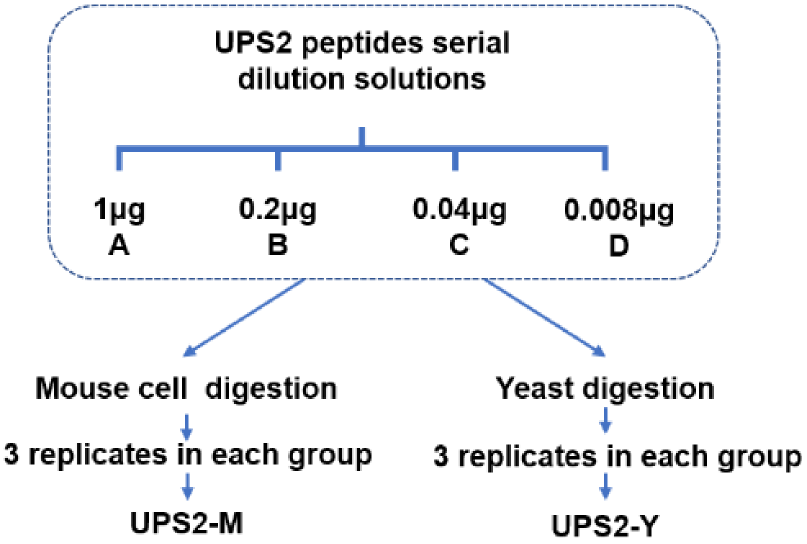


**Supplementary Fig. 3.** A series of UPS2 protein digestions (1, 0.2, 0.04, and 0.008 μg, represented as A, B, C, and D in this study) was added into an equal amount of mouse cell and yeast mixtures to build the UPS2-M and UPS2-Y datasets. This figure was modified from our previous paper (Chang et al. Anal Chem 2016, 88 (13), 6844–6851).

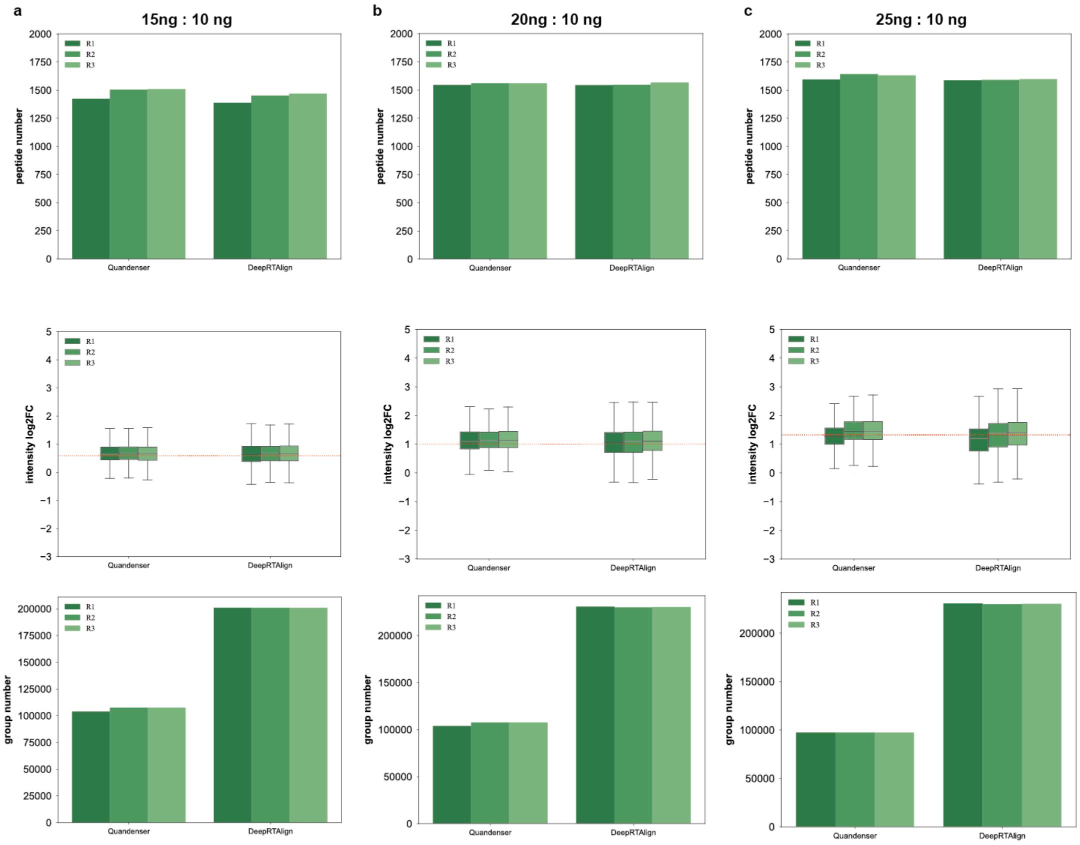


**Supplementary Fig. 4**. The number and ratio distributions of all *E. coli* peptides and the group number of aligned features between specific samples (a: 15ng/10ng, b: 20ng/10ng, and c: 25ng/10ng)

in each replicate (R1, R2 and R3) after alignment by Quandenser and DeepRTAlign. It should be noted that a group is defined as a set of aligned features in different runs.

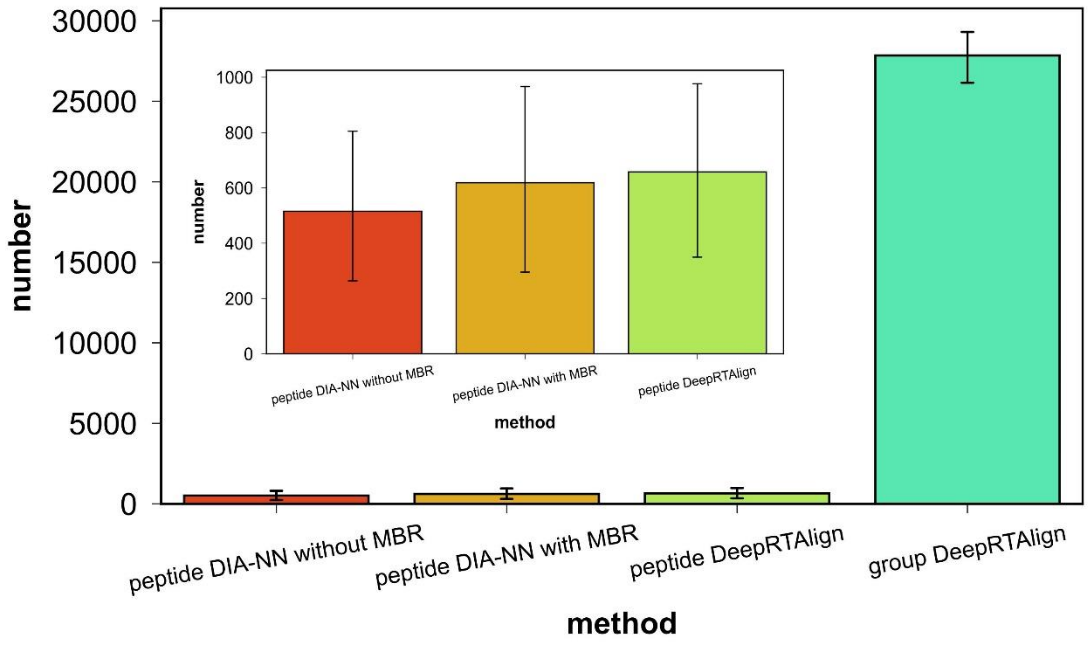


**Supplementary Fig. 5.** The peptide number and feature number of each HT22 cell. Features are extracted by Dinosaur. Only the features presented in at least two cells are considered. MBR: match between runs. Error bar indicates standard deviation. It should be noted that a group is defined as a set of aligned features in different runs.

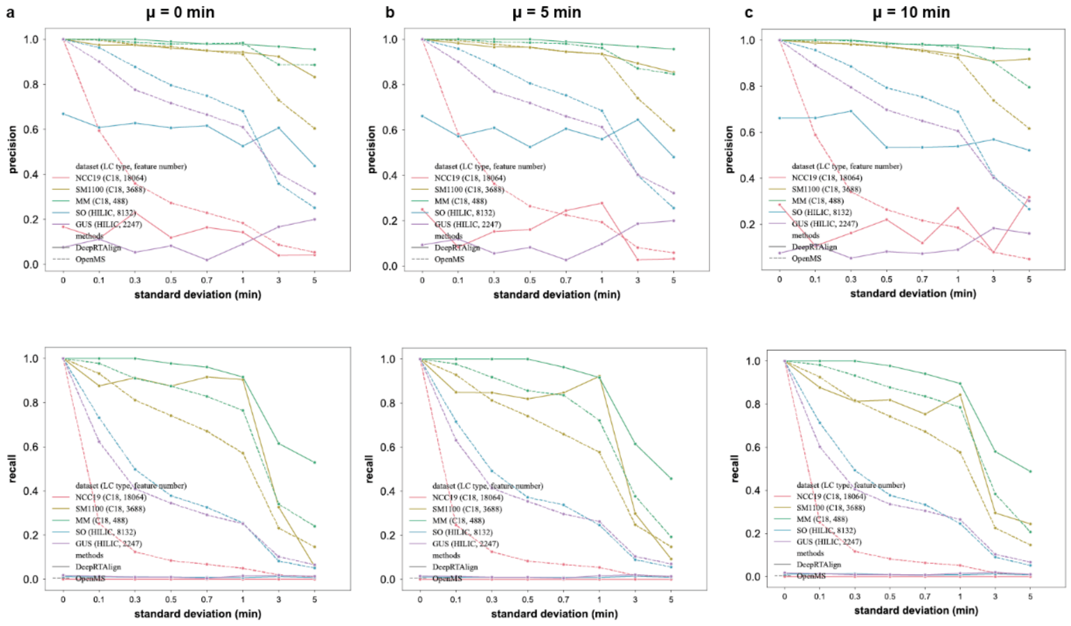


**Supplementary Fig. 6**. Comparison of DeepRTAlign and OpenMS on multiple simulated datasets generated from 5 real-world metabolomic datasets. The simulated datasets were constructed by adding normally distributed RT shifts to the corresponding real-world dataset. (a) μ=0 min. (b) μ=5

min. (c) μ=10 min. The normal distribution has an increasing σ, i.e., σ=0, 0.1, 0.3, 0.5, 0.7, 1, 3, 5 for different μ (0, 5 and 10 minutes), respectively. The FDR of DeepRTAlign's results is set to 1%.

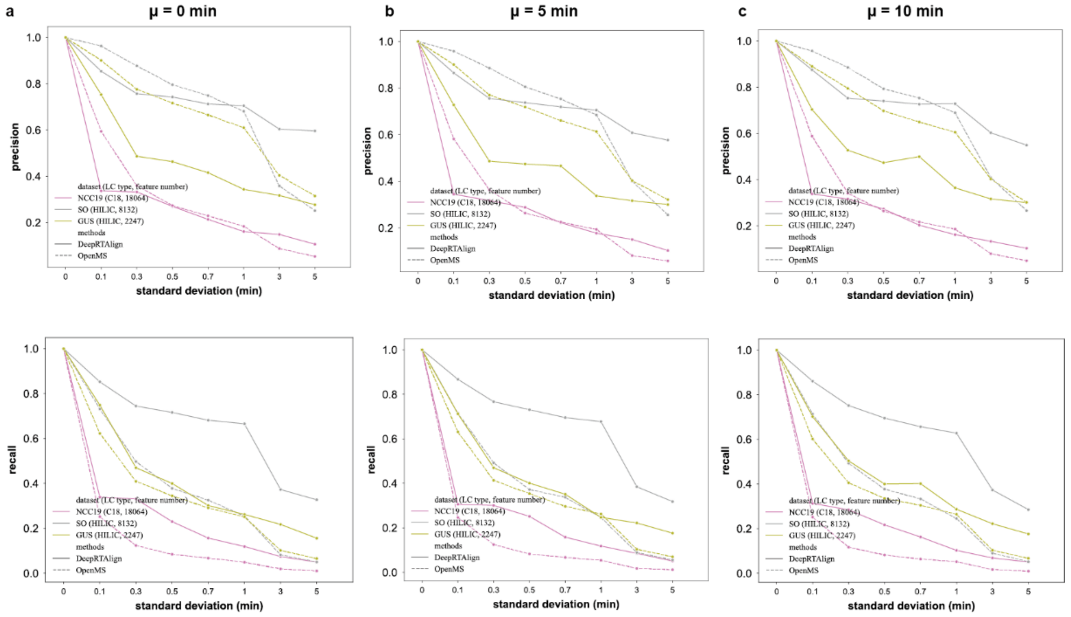


**Supplementary Fig. 7**. Comparison of DeepRTAlign and OpenMS on multiple simulated datasets generated from 3 real-world metabolomic datasets. The simulated datasets were constructed by adding normally distributed RT shifts to the corresponding real-world dataset. (a) μ=0 min. (b) μ=5 min. (c) μ=10 min. The normal distribution has an increasing σ, i.e., σ=0, 0.1, 0.3, 0.5, 0.7, 1, 3, 5 for different μ (0, 5 and 10 minutes), respectively. The FDR of DeepRTAlign's results is set to 100%.

### Supplementary Notes

#### Comparison with an in-house trained Siamese network

Aligning two samples is to compare the features within them and match the identical ones. This type of problem belongs to a field of machine learning called One-Shot Learning. Siamese network is the most commonly used architecture for this type of problem[2]. Thus, we first constructed a Siamese network for evaluation. The Siamese network has two fully-connected networks. Contrastive loss is used as the loss function. As shown in **Supplementary Table 18**, the Siamese network did not achieve the same performance of our DNN model in DeepRTAlign when using the same training set. This may be because it is hard for the Siamese network to evaluate the features that do not exist in the training set. It should be noted that the Siamese network we trained on LC-MS data was different from the network in Li et al.'s paper.
